# Sex-specific, *pdfr-1*-dependent modulation of pheromone avoidance by food abundance enables flexibility in *C. elegans* foraging behavior

**DOI:** 10.1101/2021.03.16.435685

**Authors:** Jintao Luo, Douglas S. Portman

## Abstract

To make adaptive feeding and foraging decisions, animals must integrate diverse sensory streams with multiple dimensions of internal state. In *C. elegans*, foraging and dispersal behaviors are influenced by food abundance, population density, and biological sex, but the neural and genetic mechanisms that integrate these signals are poorly understood. Here, by systematically varying food abundance, we find that chronic avoidance of the population-density pheromone ascr#3 is modulated by food thickness, such that hermaphrodites avoid ascr#3 only when food is scarce. The integration of food and pheromone signals requires the conserved neuropeptide receptor PDFR-1, as *pdfr-1* mutant hermaphrodites display strong ascr#3 avoidance even when food is abundant. Conversely, increasing PDFR-1 signaling inhibits ascr#3 aversion when food is sparse, indicating that this signal encodes information about food abundance. In both wild-type and *pdfr-1* hermaphrodites, chronic ascr#3 avoidance requires the ASI sensory neurons. In contrast, PDFR-1 acts in interneurons, suggesting that it modulates processing of the ascr#3 signal. Although a sex-shared mechanism mediates ascr#3 avoidance, food thickness modulates this behavior only in hermaphrodites, indicating that PDFR-1 signaling has distinct functions in the two sexes. Supporting the idea that this mechanism modulates foraging behavior, ascr#3 promotes ASI-dependent dispersal of hermaphrodites from food, an effect that is markedly enhanced when food is scarce. Together, these findings identify a neurogenetic mechanism that sex-specifically integrates population and food abundance, two important dimensions of environmental quality, to optimize foraging decisions. Further, they suggest that modulation of attention to sensory signals could be an ancient, conserved function of *pdfr-1*.

## INTRODUCTION

Feeding and foraging provide ideal opportunities to understand the mechanisms by which neural circuits give rise to flexible innate behaviors. Despite wide variation in the mechanics of foraging, the logic that guides decisions about whether to exploit or abandon a food resource is remarkably conserved. In animals as diverse as bees, birds, and fish, foraging decisions weigh the current benefit of feeding against the costs of exploration and the potential benefits of new food sources [1, 2]. Optimal foraging requires that animals assess not only local food abundance and quality but also other factors, such as population density, nutritional needs, and the risk of predation, to make the most advantageous decisions. Experimental studies of foraging behavior have provided important insights into the neural mechanisms underlying these calculations as well as the genetic mechanisms that have optimized them [3].

The nematode *C. elegans*, a bacterivore, feeds on microbes that colonize decaying vegetation [4]. Laboratory studies have shown that *C. elegans* modulates its feeding and exploratory behaviors in ways that are consistent with optimal foraging theory: foraging and dispersal in adults varies with food abundance [5, 6], nutritional value [7, 8], pathogenicity and toxicity [9, 10], and experience [11, 12]. Studies of wild isolates have demonstrated substantial natural variation in foraging behavior, owing in some cases to altered responses to sensory signals [6, 13–17]. Furthermore, foraging and dispersal are sexually dimorphic and sensitive to nutritional state, as solitary males will abandon a high-quality food source in search of mates, but suppress this behavior after food deprivation [18–22].

Foraging behaviors in many species are sensitive to presence of conspecifics. These can sometimes be competitors that reduce the marginal value of a crowded food source, but in other contexts, collective foraging can be more advantageous than solitary foraging [23]. There is evidence for both of these possibilities in *C. elegans*: worms will leave a food source more often if it is crowded [17], but some pheromones inhibit dispersal and exploratory behaviors [6, 16]. Further, many natural isolates feed in groups [13] and theoretical models suggest that collective foraging may be favored in patchy food environments [24]. These findings indicate that the role of population density in *C. elegans* foraging and dispersal may not be fixed. Indeed, in the wild, balancing selection can simultaneously maintain alleles that promote both strategies [16, 17].

Because of the “boom-and-bust” life cycle of *C. elegans*, in which food availability and population density can vary rapidly and dramatically [25], worms likely integrate information about these two variables when assessing the current and future quality of a food source. In *C. elegans*, population density is mainly signaled by ascarosides, derivatives of the dideoxy sugar ascarylose that serve as a modular chemical language in nematodes [26–28]. A prominent member of this group is ascr#3 (also called “C9” and “asc-ΔC9”), which, together with other ascarosides, signals population density early in larval development to influence the decision to enter the stress-induced dauer stage [29, 30]. ascr#3 is also a sex pheromone, as it is produced predominantly by adult hermaphrodites and elicits strong male-specific attraction [31–34]. In contrast, adult hermaphrodite behavioral responses to ascr#3 are typically aversive, though this depends on the context of ascr#3 presentation as well as animals’ internal state and previous experience [32, 34–38].

In our previous studies of sexual dimorphism in ascr#3 responses [33], we encountered variability in hermaphrodite ascr#3 avoidance responses, leading us to wonder whether flexible responses to population density signals might have adaptive value. Here, by systematically varying the thickness of the food lawn, we show that variability in ascr#3 avoidance can result from food abundance. We find that the conserved neuropeptide receptor PDFR-1 is necessary for this plasticity; moreover, signaling by ligand(s) produced by *pdf-1* appears to integrate information about food abundance into the ascr#3 avoidance circuit. Unexpectedly, ascr#3 avoidance in this context does not require the ADL neurons, which are known to have an important role in acute, off-food ascr#3 avoidance [34, 35], but instead depend on the ASI sensory neurons. The PDFR-1 signal likely acts downstream of ASI by modulating a distributed set of interneurons. Interestingly, we find that this modulation is sex-specific: although males possess a latent, ASI- and *pdfr-1*-dependent ascr#3 avoidance mechanism, it is not modulated by food thickness. Finally, we find that this mechanism does indeed have a potential role in worm fitness: ascr#3 can promote ASI-dependent dispersal from a food source, and this effect is markedly enhanced when food is sparse. Together, these studies identify a neurogenetic mechanism that sex-specifically integrates two distinct sensory stimuli, providing a means by which population density cues can adaptively modulate feeding and foraging decisions in the adult *C. elegans* hermaphrodite.

## RESULTS

### Hermaphrodite pheromone avoidance is modulated by food abundance

To explore the relationship between food and pheromone signals in *C. elegans* adults, we used a quadrant-format behavioral assay [33] (Figure 1A). This assay, carried out in the presence of bacterial food, allows robust quantitation of medium-term (“chronic,” 1-2 hr) attractive and aversive responses to non-volatile chemical stimuli. To control the abundance of food on the assay plate, we prepared standardized lawns of *E. coli* OP50 of three different densities (Figure 1B; see Methods for details). “T3” is a smooth, uniform lawn providing ample but not excessive food, similar to standard culture conditions. “T2” is a thin, slightly patchy lawn. “T1” is a very thin lawn with many food-free patches, mimicking a food source approaching depletion.

**Figure 1.**
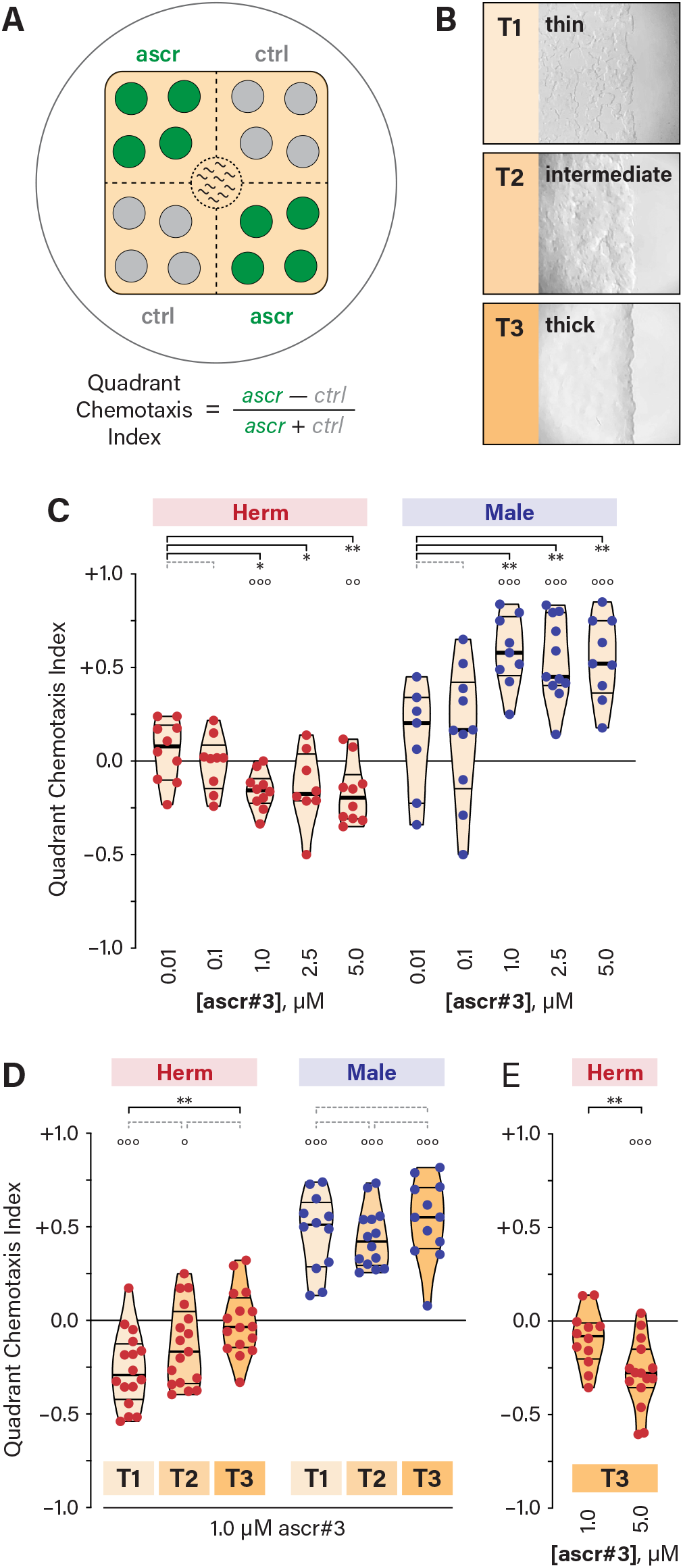
Hermaphrodite avoidance of ascr#3 depends on food abundance. **(A)** The quadrant-format ascr#3 chemotaxis assay [33]. **(B)** Photomicrographs of the edges of representative bacterial food lawns. T3, a thick food lawn; T2, a food lawn of intermediate thickness; and T1, a sparse, patchy food lawn. **(C)** Quadrant Chemotaxis Index (QCI) values for adults of both sexes in the quadrant assay in response to a range of ascr#3 concentrations in the presence of thin food (T1). **(D)** QCIs for adult of both sexes in response to 1.0 μM ascr#3 in the presence of three different food abundances (T1, T2, and T3). **(E)** QCIs for adult hermaphrodites to 1.0 and 5.0 μM ascr#3 in the presence of thick food (T3). For all quadrant assay data shown here and in the following figures, data points are colored by sex (hermaphrodites in red and males in blue) and represent the QCI from a single assay containing *n* = 10 worms. Violin plots illustrate the distribution of the data and are shaded according to food thickness (T1 = lighter; T3 = darker). The median and 25%/75% quartile intervals are indicated by thick and thin black lines, respectively. Statistical comparisons between groups are indicated with black brackets and asterisks (* *p* ≤ 0.05; ** *p* ≤ 0.005; *** *p* ≤ 0.001) or dotted gray brackets (*p* > 0.05). Open circles above each column of data indicate the results of one-sample *t-*tests, carried out to determine whether the observed QCI differs significantly from zero (° *p* ≤ 0.05; °° *p* ≤ 0.005; °°° *p* ≤ 0.001).

We first examined behavioral responses to ascr#3 under thin food conditions (Figure 1C and S1). In hermaphrodites, we observed modest but consistent avoidance of 1–5 μM ascr#3, but no response to lower concentrations (0.1 and 0.01 μM) was seen. In males, 1–5 μM ascr#3 elicited robust attraction, but there was little or no response to lower concentrations (0.01 μM and 0.1 μM). Thus, as expected, hermaphrodites and males exhibit qualitatively distinct responses to ascr#3 when assayed with sparse food, and the concentration-dependence of these responses was similar.

Next, we examined ascr#3-evoked behavior under different food-abundance conditions. In adult hermaphrodites, consistent ascr#3 avoidance was apparent in the context of thin lawns (T1 and T2), but, surprisingly, we found that animals were indifferent to this stimulus in the presence of thicker T3 food (Figure 1D). In contrast, adult males displayed a robust preference for the ascr#3-containing quadrants under all three conditions, with no apparent effect of food thickness (Figure 1D). Thus, food thickness modulates the intensity of ascr#3 aversion in hermaphrodites but has no apparent effect on ascr#3 attraction in males.

The modulation of ascr#3 aversion by food abundance could reflect active integration of food and pheromone signals by the hermaphrodite nervous system, with the appetitive value of thick food outweighing the aversive effects of ascr#3. Alternatively, thick food might simply block ascr#3 detection or response. We found that animals exhibited avoidance of more concentrated ascr#3 (5 μM) on thick food (Figure 1E), consistent with the idea that the hermaphrodite nervous system actively integrates the opposing signals of food abundance and pheromone to influence navigation.

### PDF neuropeptide signaling modulates pheromone avoidance

We next sought to identify the mechanisms underlying food-dependent plasticity in ascr#3 avoidance. Because the conserved neuropeptide receptor PDFR-1 [39] has previously been associated with both food-dependent and sex-specific behavioral states [19, 40], we examined the behavior of *pdfr-1* mutants. Interestingly, on thin food, *pdfr-1* null mutant hermaphrodites displayed markedly enhanced ascr#3 avoidance compared to controls (Figure 2A, S2A). Single null mutations in *pdf-1* and *pdf-2*, which encode ligands for PDFR-1 [39], caused little change in ascr#3 avoidance. However, we observed enhanced avoidance in *pdf-1; pdf-2* double mutants (Figure 2A), suggesting functional redundancy between these ligands. We detected no significant difference between the ascr#3 avoidance behavior of *pdfr-1* and *pdf-1; pdf-2* mutants, but we cannot rule out the possibility that additional ligands contribute to the regulation of PDFR-1 activity.

**Figure 2.**
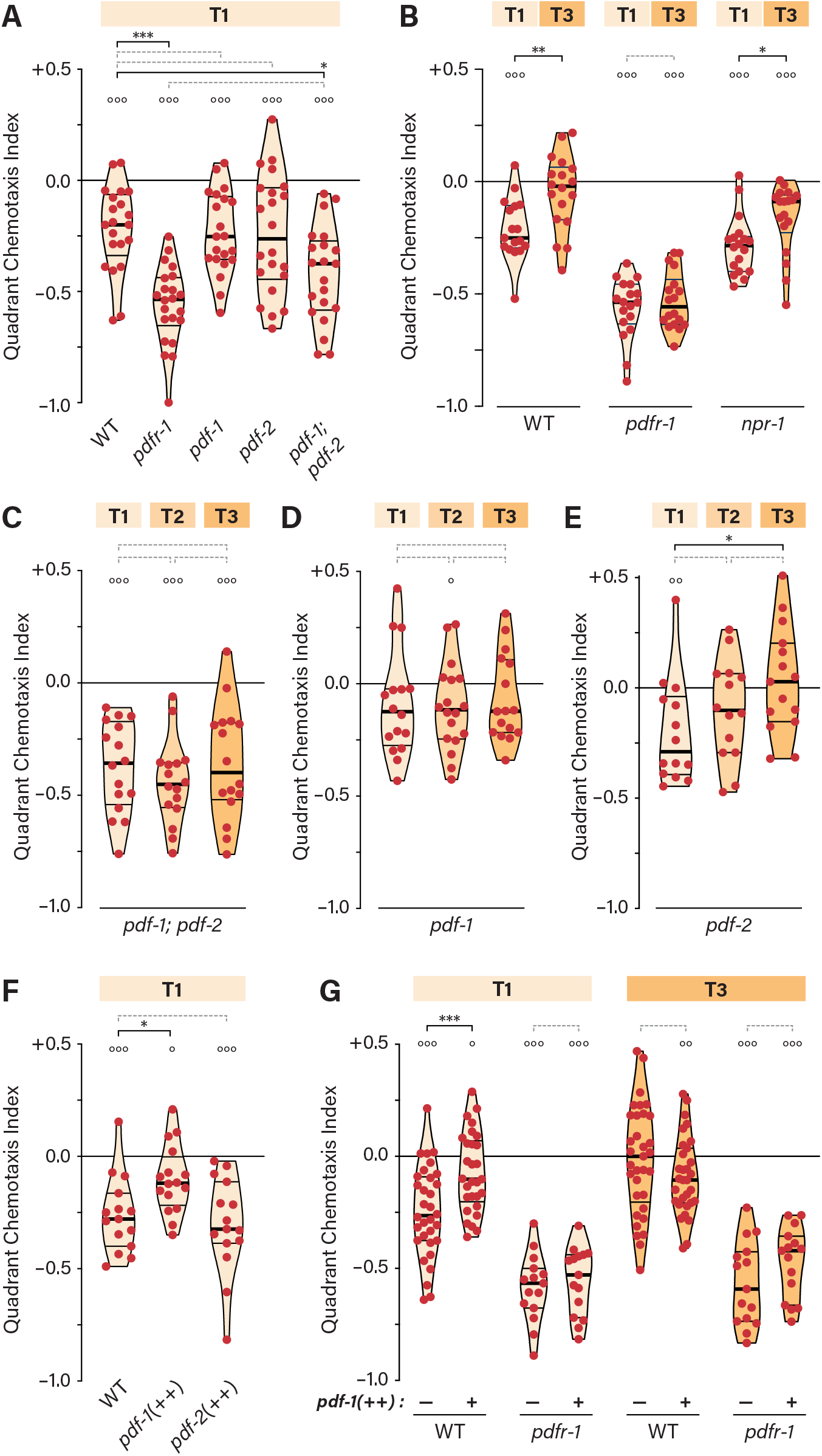
PDFR-1 signaling couples food abundance to the repression of ascr#3 avoidance. **(A)** QCIs for WT, *pdfr-1*, *pdf-1*, *pdf-2*, and *pdf-1; pdf-*2 mutant hermaphrodites with T1 food. **(B)** QCIs for WT, *pdfr-1*, and *npr-1* hermaphrodites with T1 and T3 food. **(C-E)** QCIs for *pdf-1*; *pdf-2* (C), *pdf-1* (D), and *pdf-2* (E) mutant hermaphrodites with three different food conditions. **(F)** QCIs for WT hermaphrodites and hermaphrodites overexpressing *pdf-1* (*pdf-1(++)*) and *pdf-2* (*pdf-2(++)*) with T1 food. **(G)** QCIs for WT, *pdf-1(++)*, *pdfr-1*, and *pdfr-1; pdf-1(++)* hermaphrodites with T1 and T3 food. See legend to Figure 1 for details of statistical analyses.

Because larval exposure to ascr#3 can alter adult ascr#3 responses [37], our results could point to a role for *pdfr-1* in this experience-dependent plasticity. However, we found that the ascr#3 avoidance measured by the quadrant assay was independent of previous ascr#3 exposure, as the loss of *daf-22*, an enzyme necessary for synthesis of ascr#3 and other short-chain ascarosides [41], did not change the behavior of WT or *pdfr-1* hermaphrodites (Figure S2B).

### PDFR-1 signaling intensity likely encodes food thickness information

Interestingly, we found that the ascr#3 responses of *pdfr-1* mutants were insensitive to food thickness: *pdfr-1* hermaphrodites showed strong aversion to ascr#3 regardless of food abundance, with no apparent difference between thin and thick food conditions (Figure 2B). Because responses to ascr#3 are also modulated by the neuropeptide receptor *npr-1* [34, 35], we examined the behavior of *npr-1* mutants in different food contexts. On thin food, *npr-1* hermaphrodites displayed ascr#3 avoidance comparable to WT; furthermore, ascr#3 avoidance remained sensitive to food abundance, as it was significantly blunted in the presence of thick food (Figure 2B). Thus, *pdfr-1*, but not *npr-1*, is required for modulation of ascr#3 aversion by food abundance.

Consistent with this requirement for *pdfr-1*, the food-dependence of ascr#3 avoidance was eliminated in *pdf-1; pdf-2* double mutants (Figure 2C). Surprisingly, food-dependence was also absent in *pdf-1* single mutants (Figure 2D), but it remained intact in *pdf-2* mutants (Figure 2E). Thus, although ligands encoded by both genes appear to have roles in the modulation of ascr#3 aversion, *pdf-1* has a specific role in linking food thickness to the strength of the aversive response.

We noted that the difference in ascr#3 avoidance between WT and *pdfr-1* hermaphrodites appeared to be greater on thick food than on thin food (Figure 2B). This suggests that the intensity of PDFR-1 signaling in WT animals might encode information about food thickness. To test this, we asked whether an increase in *pdfr-1* signaling would be sufficient to cause the behavior of animals assayed on thin food to be more like that typically seen on thick food. To bring this about, we used previously validated transgenes carrying extra copies of wild-type *pdf-1* or *pdf-2* [39]. Consistent with our hypothesis, we found that *pdf-1* overexpression reduced ascr#3 avoidance on thin food (Figures 2F, G). Overexpression of *pdf-2*, in contrast, had no apparent effect (Figure 2G), suggesting that *pdf-1*, but perhaps not *pdf-2*, conveys food thickness information. In agreement with this possibility, overexpression of *pdf-1* had no apparent effect on behavior in the presence of thick food (Figure 2G). As expected, *pdfr-1* was required for the effect of *pdf-1* overexpression, as *pdfr-1; Ex[pdf-1(++)]* animals behaved comparably to *pdfr-1* mutants in both food conditions tested (Figure 2G). These findings, together with the loss-of-function data described above, indicate that the intensity of PDFR-1 signaling, driven primarily by *pdf-1* activity, provides an instructive internal representation of food thickness that modulates ascr#3 avoidance.

### The ASI sensory neurons are an essential component of the ascr#3 avoidance circuit

To better understand how *pdfr-1* modulates ascr#3 avoidance, we sought to characterize the neuronal requirements for this behavior. Previous studies have found that acute aversive and attractive responses to ascr#3 in the absence of food are mediated by the sensory neurons ADL and ASK, respectively [32, 34, 35, 42]. However, we found that genetic ablation of neither of these pairs had an apparent effect on the chronic ascr#3 avoidance measured by the quadrant assay, either in wild-type or *pdfr-1* hermaphrodites (Figures 3A, B). Thus, the longer-term, on-food responses we examine here are likely mediated by mechanisms distinct from those controlling acute ascr#3 responses in the absence of food. This is reminiscent of our previous finding that male ascr#3 attraction in the quadrant assay depends primarily on ADF [33], rather than on ASK or the male-specific CEM neurons, both of which are important for ascr#3 attraction in other contexts [32, 35].

**Figure 3.**
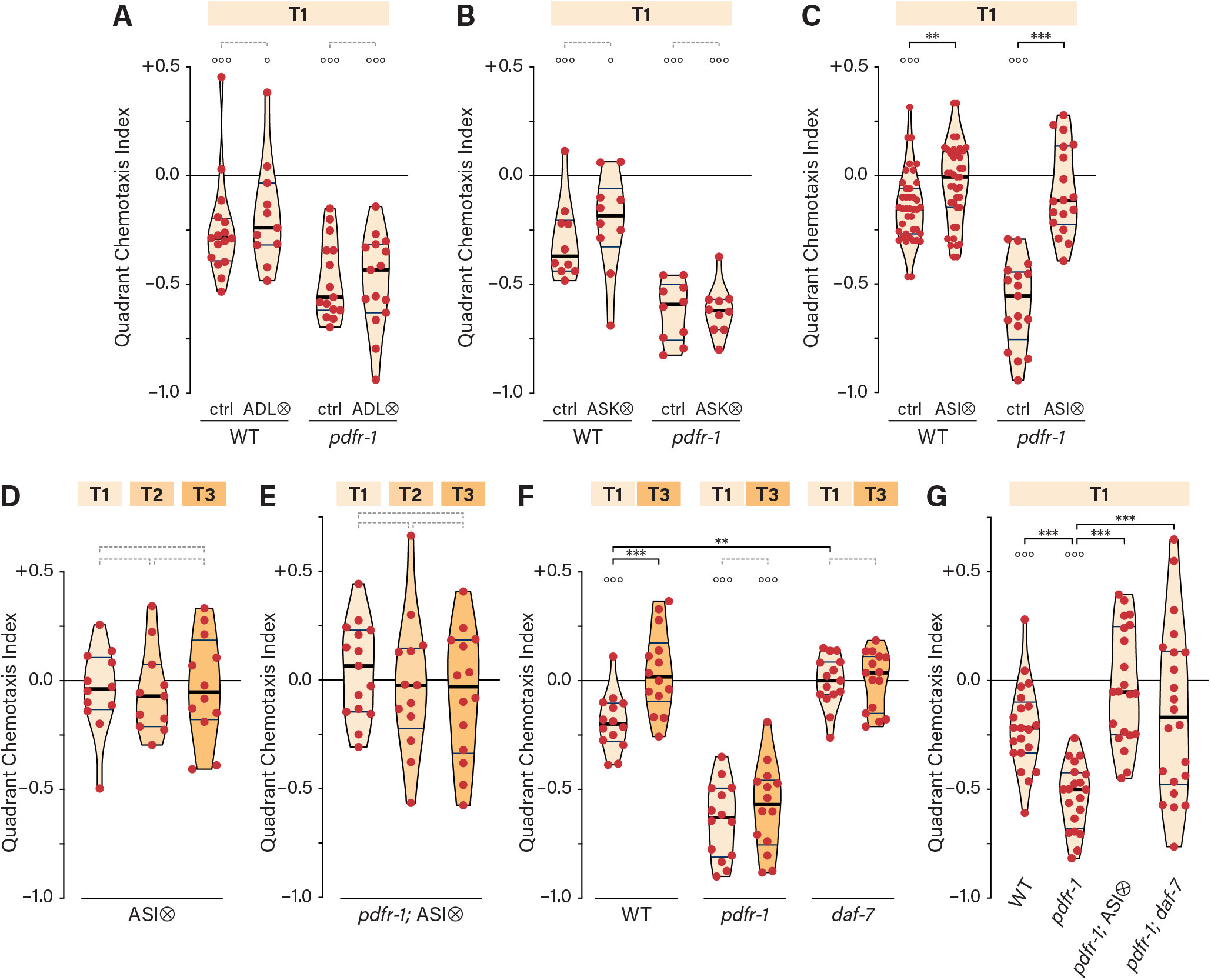
The ASI neurons mediate *pdfr-1*-dependent ascr#3 avoidance in hermaphrodites. **(A, B)** QCIs for control and ADL-ablated (ADL⊗) (A) or ASK-ablated (ASK⊗) (B) hermaphrodites in a wild-type or *pdfr-1* background with T1 food. **(C)** QCIs for WT and ASI-ablated (ASI⊗) hermaphrodites in a wild-type or *pdfr-1* background with T1 food. **(D, E)** QCIs for ASI⊗ (D) or *pdfr-1*; ASI⊗ (E) hermaphrodites with three food conditions. **(F)** QCIs for WT, *pdfr-1*, and *daf-7* hermaphrodites with T1 and T3 food. **(G)** QCIs for WT, *pdfr-1*, *pdfr-1*; ASI⊗ and *pdfr-1; daf-7* hermaphrodites with T1 food. See legend to Figure 1 for details of statistical analyses.

We next considered the ASI neurons, which can detect ascarosides in other contexts, including the decision to enter the stress-resistant dauer stage [43] and the response to the exploration-suppressing pheromone icas#8 [16, 44]. We found ascr#3 avoidance was abolished upon ASI ablation, both in wild-type and *pdfr-1* backgrounds (Figures 3C), regardless of food thickness (Figures 3D, E). The requirement for ASI in both wild-type and *pdfr-1* mutants indicates that the strong ascr#3 avoidance of *pdfr-1* mutants likely results from increased activity of the same mechanism that generates avoidance in wild-type hermaphrodites.

These results suggested that ASI might directly detect ascr#3. However, using validated GCaMP transgenes and a range of pheromone concentrations, others have seen no evidence of ascr#3-evoked calcium responses in ASI (E. DiLoreto and J. Srinivasan, pers. comm.; K. Kim and P. Sengupta, pers. comm.). This is consistent with previous findings that ascarosides can engage signaling mechanisms that act over longer timescales and are not accompanied by calcium transients [15, 30, 44]. The simplest interpretation of our results is that ASI directly detects ascr#3; however, it is also possible that ASI is required downstream of ascr#3 detection or in parallel with it.

We also considered the possibility that ASI detects or implements the thin-food state, since ASI-ablated hermaphrodites mimic thick-food behavior even in the thin-food environment (Figure 3C). However, this appeared not to be the case, as ASI ablation in *pdfr-1* mutants eliminated pheromone avoidance on thin food, rather than recapitulating the moderate ascr#3 repulsion typical of *pdfr-1* hermaphrodites on thick food (Figure 3C).

To understand the role of ASI in ascr#3 aversion, we examined the TGFβ-superfamily ligand DAF-7 [45, 46], one of several neuromodulators produced by ASI. Interestingly, we found that *daf-7* mutant hermaphrodites exhibited no response to ascr#3 on either thin or thick food, reminiscent of ASI-ablated hermaphrodites (Figure 3F). Further, *daf-7* suppressed the enhanced ascr#3 avoidance of *pdfr-1* hermaphrodites on thin food (Figure 3G). Although the behavioral variability of *pdfr-1; daf-7* double mutants was exceptionally high, these results indicate that the function of ASI in ascr#3 avoidance might be mediated by *daf-7* and that PDFR-1 might modulate the neural circuit that the *daf-7* signal engages.

### PDFR-1 functions in multiple interneurons to modulate ascr#3 avoidance

We next sought to identify the site of action of *pdfr-1*. Previous work has shown that *pdfr-1* is expressed in multiple head and tail neurons, as well as peripheral tissues such as body wall muscle [19, 39, 47]. Using a previously described intersectional strategy [40], we tested the ability of neuron-specific nCre constructs, together with a conditionally activatable floxed *Ppdfr-1::Inv[PDFR-1::SL2::GFP]* transgene, to rescue the enhanced ascr#3 avoidance phenotype of *pdfr-1* hermaphrodites. This transgene produces functional *pdfr-1* and *GFP* transcripts only in neurons in which both the *Ppdfr-1* promoter and nCre are active (Figure 4A).

**Figure 4.**
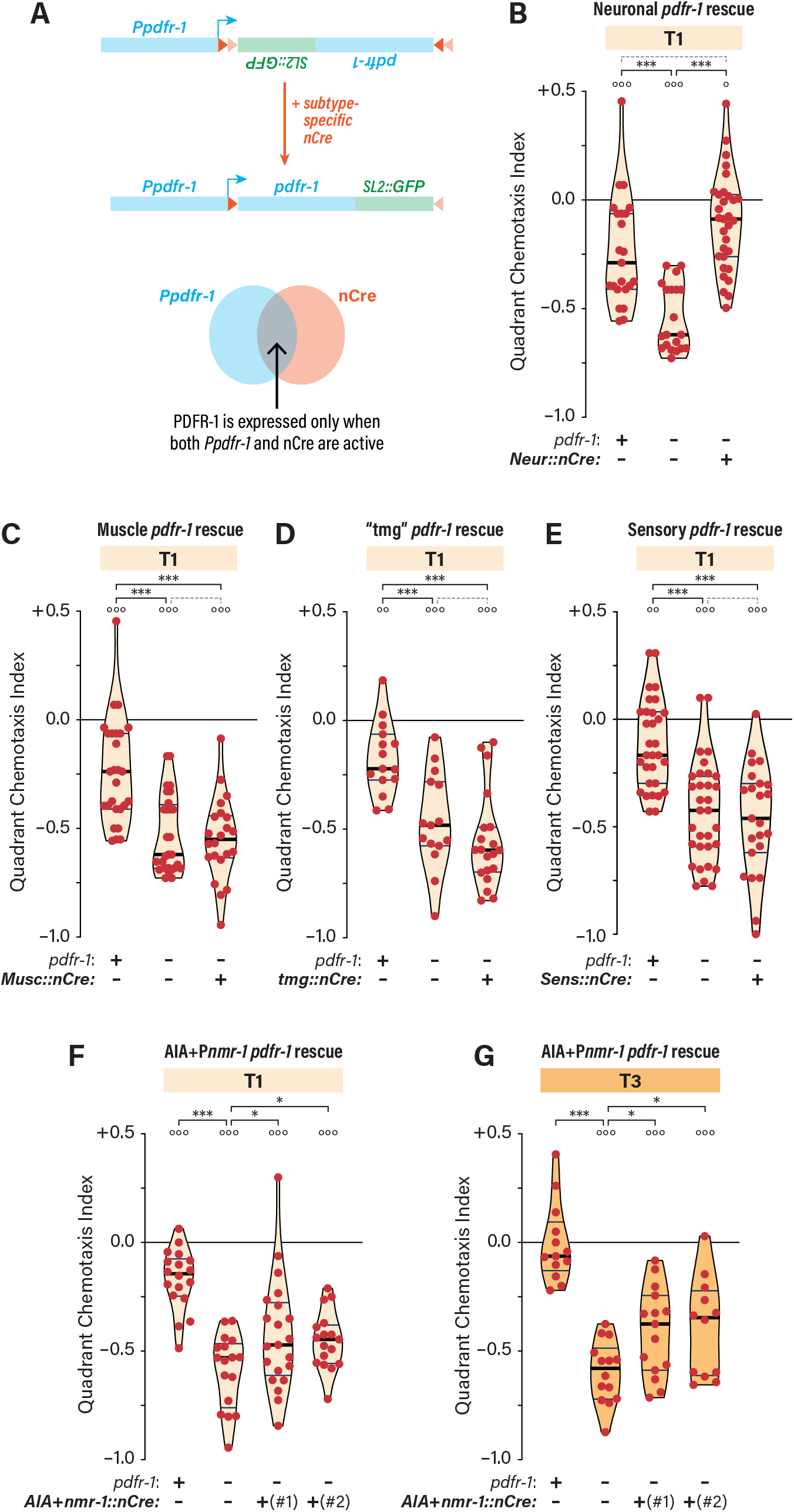
*pdfr-1* acts in a distributed set of interneurons to repress ascr#3 avoidance in hermaphrodites. **(A)** The strategy for Cre-based conditional rescue of *pdfr-1*, developed previously [40]. **(B-G)** QCIs for hermaphrodites carrying the conditional *pdfr-1* transgene *kyEx4648* with the indicated *pdfr-1* genotypes at the endogenous *pdfr-1* locus (+, wild-type; –, mutant) without or with transgenes driving nCre expression pan-neuronally (“ *Neur::nCre*”) (B), in muscle (“ *Musc::nCre*”) (C), in a previously described set of interneurons in which *pdfr-1* functions to regulate motor state [40] (“ *tmg::nCre*”) (D), in sensory neurons (“ *Sens::nCre*”) (E), and in AIA and neurons expressing *nmr-1* (“ *AIA+nmr-1::nCre*”) (F, G), with T1 (B-F) or T3 (G) food. See legend to Figure 1 for details of statistical analyses.

Consistent with the idea that *pdfr-1* acts in the nervous system, pan-neural nCre expression was sufficient to completely rescue the *pdfr-1* defect, but muscle expression of nCre was not (Figures 4B, C). Next, we tested a combination of three nCre transgenes (“ *tmg::nCre*,” driven by *Ptdc-1*, *Pmod-1*, and *Pglr-3*) that was previously shown to provide partial rescue of the persistent dwelling behavior of *pdfr-1* mutants [40]. However, we observed no rescue of the ascr#3 avoidance phenotype with this transgene (Figure 4D). To ask whether *pdfr-1* functions in sensory neurons, we used *Posm-6::nCre*, but again saw no rescuing activity (Figure 4E). This suggests that *pdfr-1* is unlikely to function in ASI, an interpretation strengthened by the absence of a GFP signal in animals carrying an ASI/AWA-specific *Pgpa-4::nCre* transgene together with the conditional *pdfr-1* transgene (data not shown).

We next generated and tested several other cell-type-specific nCre constructs in combination with the conditional *pdfr-1* transgene. These strains either exhibited no detectable GFP (*Pmec-10,* touch neurons) or failed to rescue the ascr#3 avoidance phenotype (*Pmod-1 + Posm-6,* GFP seen in several sensory neurons and interneurons; *Ptdc-1 + Pglr-3 + Pnmr-1,* GFP in many interneurons; and *Pgcy-28.d,* GFP in AIA interneurons) (data not shown). However, with a different combination, *Pgcy-28.d::nCre* + *Pnmr-1::nCre*, we observed partial but consistent and significant reduction of the enhanced ascr#3 avoidance of *pdfr-1* mutants in both thin-food and thick-food conditions (Figures 4F, G). In these animals, GFP was detected in several interneurons, including AIA, PVC, and one or more *nmr-1*-expressing head interneurons (AVA, AVD, AVE, AVG, and RIM). From these results, we conclude that PDFR-1 does not have one discrete site of action. Rather, its distributed role suggests that PDFR-1 coordinately modulates multiple components of the ascr#3 avoidance circuit in response to PDF neuropeptides.

### The modulatory effects of food thickness are sex-specific

Males and hermaphrodites have markedly different responses to ascr#3 [32]. In males, sex-specific detection of ascr#3 by the sex-shared ADF neurons promotes attraction in the quadrant assay [33]. Consistent with this, we found that *pdfr-1* and ASI are dispensable for male ascr#3 attraction (Figures 5A, B).

**Figure 5.**
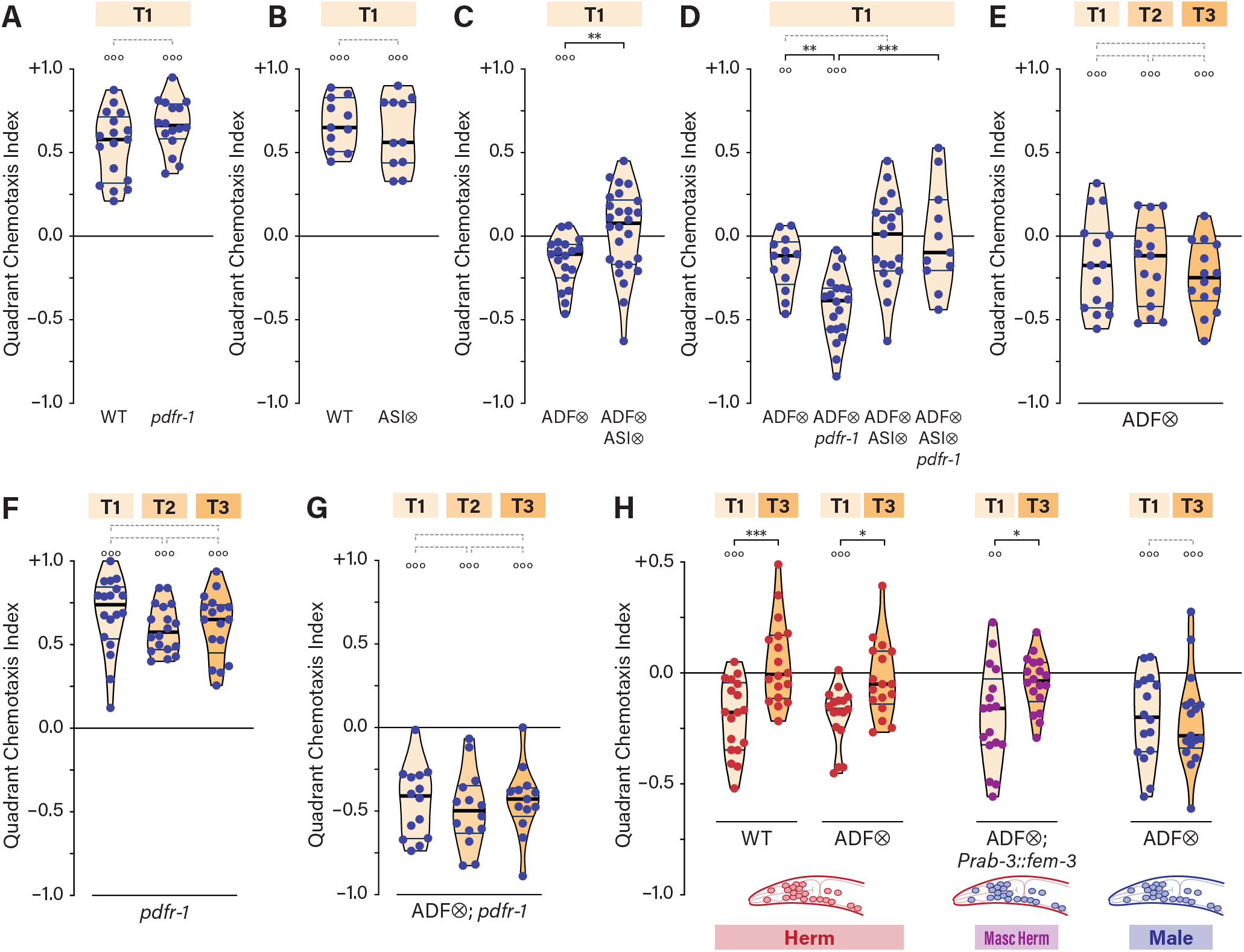
A latent ASI-dependent ascr#3 avoidance mechanism in males is independent of food abundance. **(A, B)** QCIs for WT and *pdfr-1* (A) or ASI⊗ (B) males with T1 food. **(C)** QCIs for ADF⊗ and ADF⊗; ASI⊗ males with T1 food. **(D)** QCIs for ADF⊗, ADF⊗; *pdfr-1*, ADF⊗; ASI⊗, and *pdfr-1*; ADF⊗; ASI⊗ males with T1 food. **(E-G)** QCIs for ADF⊗ (E), *pdfr-1* (F), and ADF⊗; *pdfr-1* (G) males with T1, T2, and T3 food. **(H)** QCIs for WT hermaphrodites, ADF⊗ hermaphrodites, ADF⊗ hermaphrodites carrying the pan-neuronal masculinization transgene *Prab-3::fem-3*, and ADF⊗ males with T1 and T3 food. See legend to Figure 1 for details of statistical analyses.

When the influence of ADF is removed by genetic feminization or ablation, males display aversive responses to ascr#3, similar to those of wild-type hermaphrodites [33]. Therefore, we asked whether this underlying aversive drive is mediated by a sex-shared mechanism. Consistent with this, we found that simultaneous ablation of ADF and ASI ablation eliminated all response to ascr#3 in males (Figure 5C). Further, loss of *pdfr-1* enhanced the ascr#3 avoidance of ADF-ablated males, and this enhanced avoidance also required ASI (Figure 5D). Together, these findings indicate that the ascr#3 avoidance behavior revealed by ADF ablation is mediated by a mechanism shared by both sexes.

Despite this common mechanism, however, we found that the behavior of ADF-ablated males was not completely equivalent to that of hermaphrodites. In particular, even in the presence of thick food, ADF-ablated males continued to manifest clear aversion to ascr#3 (Figure 5E). Further, we observed no effect of food thickness on ascr#3 attraction in *pdfr-1* males or on ascr#3 aversion in ADF-ablated *pdfr-1* males (Figures 5F, G). Thus, under these conditions, the modulation of ascr#3 avoidance by food thickness is a hermaphrodite-specific feature of the nervous system.

One possible explanation for this sex-specificity is that genetic sex might functionally configure shared circuitry, such that the integration of food thickness and ascr#3 signals is enabled only in hermaphrodites. Alternatively, sex-specific components of the nervous system or a signal from non-neuronal tissues might bring about this sex difference. To investigate this, we examined hermaphrodites in which the nervous system was genetically masculinized by pan-neuronal expression of the male sexual regulator *fem-3* [48]. This manipulation has previously been shown to functionally sex-reverse many behaviors mediated by shared circuits, including chemosensation and locomotion [22, 33, 49–51]. To prevent masculinized ADF neurons from driving ascr#3 attraction in these hermaphrodites [33], we examined ADF-ablated *Prab-3::fem-3* hermaphrodites. Interestingly, pan-neuronal masculinization did not disrupt the modulatory effects of food thickness: thick food eliminated ascr#3 aversion in these animals, just as it does in WT hermaphrodites (Figure 5H). This suggests a role for sex-specific neurons and/or non-neuronal tissues in programming the nervous system’s integration of food and pheromone signals.

### Foraging decisions that integrate food abundance and population density cues require ASI

Finally, we explored the potential ecological significance of food- and *pdfr-1*-dependent modulation of ascr#3 avoidance. In particular, we wondered whether this plasticity might allow hermaphrodites to incorporate population density into their assessment of the value of an existing food resource. When food is abundant, it might be adaptive for hermaphrodites to disregard population density, allowing them to rapidly exploit a plentiful resource. When food is scarce, however, population density might have higher salience, as the amount of competition for a very limited resource could be a key predictor of the future potential of a food patch.

To explore this, we measured the propensity of wild-type and ASI-ablated hermaphrodites to disperse from food sources of variable quality. We designed an assay in which a thin (T1) or thick (T3) food source, supplemented with vehicle control or 1 μM ascr#3, is placed at the center of a 10 cm agar plate (Figure 6A). Surrounding this, but separated from the source patch by 1.8 cm, are four rectangular patches of thick food without ascr#3, providing favorable alternative environments. At the beginning of the assay, 50 young adult hermaphrodites (25 wild-type and 25 ASI-ablated, distinguishable by a fluorescent marker) were deposited on the source patch. After 3 hr, we scored the number of animals of each genotype that had migrated to the peripheral thick food lawns.

**Figure 6.**
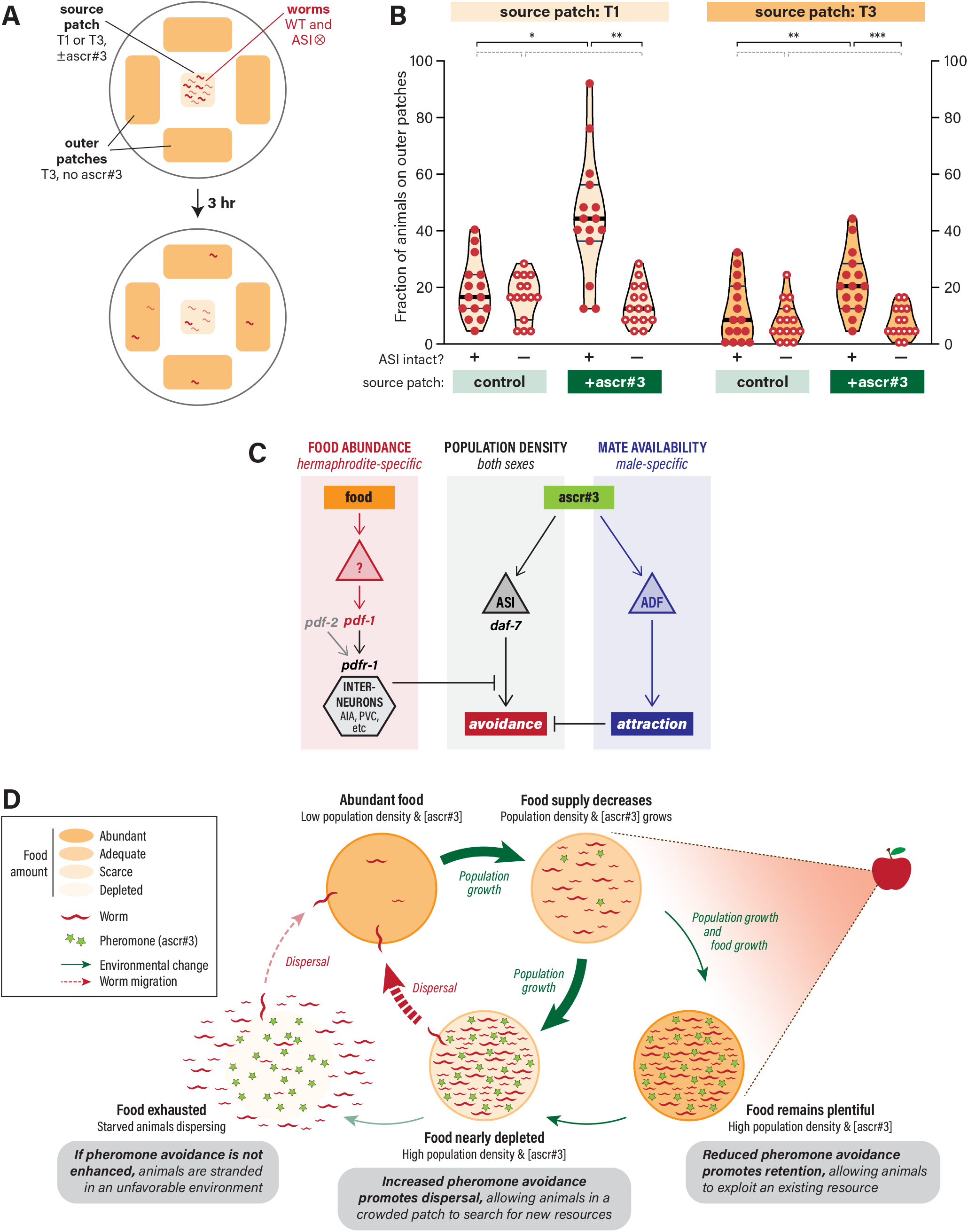
Modulation of ascr#3 aversion by food thickness enables context-dependent flexibility in *C. elegans* foraging decisions. **(A)** The behavioral assay used to measure the effects of ascr#3 and food thickness on foraging decisions. See text for details. **(B)** Foraging rates, shown as the frequency of animals dispersing to the outer food sources, of WT and ASI⊗ hermaphrodites when placed on a source patch of thick (T3) or thin (T1) food, without (–) or with (+) ascr#3. Each data points represents a single genotype in a single assay containing *n* = 25 worms per genotype. Filled and open circles indicate control and ASI-ablated animals, respectively. Violin plots illustrate the distribution of the data and are shaded according to food thickness at the source patch (T1 = lighter; T3 = darker). The median and 25%/75% quartile intervals are indicated by thick and thin black lines, respectively. Statistical comparisons between groups are indicated with black brackets and asterisks (* *p* ≤ 0.05; ** *p* ≤ 0.005; *** *p* ≤ 0.001) or dotted gray brackets (*p* > 0.05). **(C)** A neural circuit model showing the parallel inputs of food abundance, ascr#3 via ASI (a non-sex-specific input that population density), and ascr#3 via ADF (a male-specific input that mate availability). Hermaphrodite- and male-specific aspects are shown in red and blue, respectively. **(D)** A proposed model illustrating the adaptive value of context-dependent flexibility in *C. elegans* foraging behavior based on the integration of food-thickness and population density information via ascr#3. See text for details.

When the source was a patch of thin food, we found that relatively few animals migrated out of it, regardless of whether ASI had been ablated (Figure 6B). However, supplementing the source patch with ascr#3 brought about a marked increase in dispersal rate, with nearly 50% of animals migrating to the outer patches on average (Figure 6B). When ASI was ablated, however, addition of ascr#3 did not increase dispersal rates above baseline (Figure 6B). Thus, consistent with our findings in the quadrant assay, ascr#3 serves as a potent driver of dispersal when food is scarce. Furthermore, the effects of ascr#3 are completely dependent on the presence of ASI.

When the source patch was thick food, we again observed relatively few animals dispersing (Figure 6B). Adding ascr#3 to a thick-food source patch again caused an increase in dispersal, but the magnitude of this effect was far smaller than on thin food (Figure 6B). Furthermore, the effect of ascr#3 on dispersal was eliminated by ASI ablation. Thus, even on thick food, ascr#3 can be perceived as an aversive cue through an ASI-dependent mechanism. However, the ability of ascr#3 to promote dispersal from a food source is markedly blunted when food is abundant.

Together, these results show that flexibility in ascr#3 aversion allows hermaphrodites to modulate feeding and foraging decisions according to population density and food abundance, two key determinants of the quality of a food patch. The ASI-dependence of this plasticity strongly suggests that it arises through the same *pdfr-1*-dependent integration mechanism that modulates ascr#3 aversion in the quadrant assay. Further, because it provides a means for animals to optimize their dispersal from a food source, this mechanism likely has adaptive value in the wild.

## DISCUSSION

Foraging decisions are central to an animal’s survival. In the presence of a food source, animals must weigh the value of continuing to exploit this resource against the risks and benefits of abandoning it. Guided by genetically specified rules, this calculation must consider both the quality and amount of existing food as well as its rate of depletion, a primary determinant of which is local population density. Previous studies have demonstrated that all of these factors can modulate the propensity of *C. elegans* hermaphrodites to disperse from an existing resource [5, 6, 17]. Here, we show that the *C. elegans* nervous system actively integrates information about food abundance and population density to modulate behavioral decisions. Further, we identify a mechanism underlying this integration and find that it enables plasticity in feeding and foraging behaviors. Thus, our results provide new insight into the neural mechanisms that implement context-dependent behavioral choices in *C. elegans* and the genetically specified rules by which these mechanisms operate.

In our proposed mechanism (Figure 6C), two parallel sensory inputs of food thickness and ascr#3 concentration allow hermaphrodites to modulate the behavioral response to pheromones according to food thickness. When food is sparse, ascr#3 avoidance is manifested, but when food is more abundant, ascr#3 avoidance is suppressed. Thus, population density becomes particularly influential only when food depletion is imminent. This may allow hermaphrodites to disperse from a crowded, rapidly depleting food source before it is completely exhausted, avoiding potential starvation (Figure 6D). When food is abundant, however, population density is less relevant, likely allowing animals to continue to exploit an existing resource. Thus, based on environmental conditions, the hermaphrodite nervous system assigns a weight to population density in its calculations about the future potential of a given food source.

In the context of the on-food behavioral assay used here, we find that ascr#3 is detected through at least two distinct sensory mechanisms. Importantly, the circuits mediating ascr#3 responses appear to be context-specific, as the behaviors elicited by ascr#3 in other settings (*e.g.*, the acute drop test) have distinct neuronal and genetic requirements [32, 34, 35]. Here, one of the two ascr#3 inputs depends on the ASI sensory neurons and drives an aversive response. While we found no indication that ASI exhibits a calcium response to ascr#3 stimulation, previous work indicates that several classes of sensory neurons in *C. elegans*, including ASI, can detect ascaroside pheromones directly without displaying a calcium transient [15, 30, 44]. Interestingly, these responses seem to act over a longer timescale than typical calcium-based neuronal signals, suggesting that they mediate “primer” effects of pheromones [16]; this is consistent with the 2-hr timescale of our quadrant assay. Thus, we favor the idea that the aversive drive elicited by ascr#3 is signaled directly by ASI, but our results are also consistent with the possibility that ASI is required for the effects of an ascr#3 signal from other sensory neuron(s). We find that ASI’s function depends on the TGFβ-superfamily signal DAF-7, which, under typical conditions, is produced in hermaphrodites predominantly by ASI [45]. Whether the DAF-7 signal acts instructively or permissively with respect to ascr#3 detection will be an important question for future research.

Our results indicate that a separate sensory stream, acting in parallel with these ascr#3 inputs, signals food abundance. While we do not yet know where this information originates, our findings strongly suggest that it is represented internally by the level of activity of the neuropeptide receptor PDFR-1. Multiple lines of evidence support this idea. Animals lacking *pdfr-1* function, or the function of the two ligand-encoding genes *pdf-1* and *pdf-2*, are completely insensitive to the modulatory effects of food thickness. These animals display constitutively strong ascr#3 aversion that depends on ASI. Because this aversion is even stronger than that seen under thin-food (T1) conditions, we infer that low levels of PDFR-1 signaling are active even when animals are feeding on thin food, slightly blunting the aversive response. Under thick-food (T3) conditions, *pdfr-1* and *pdf-1; pdf-2* hermaphrodites still strongly avoid ascr#3, even though wild-type hermaphrodites are essentially indifferent to it. This indicates that when wild-type animals are feeding on thick food, high levels of PDFR-1 signaling completely inhibit ascr#3 aversion. Our results are most consistent with the idea that ligands produced by both *pdf-1* and *pdf-2* are involved in this signal, but that *pdf-1*-derived ligand(s) likely have a more important role. Importantly, increasing PDFR-1 signaling by overexpressing of *pdf-1* in animals feeding on thin food is sufficient to cause them to behave as though they were in the presence of thick food—that is, they are now indifferent to ascr#3—but overexpressing of *pdf-1* in animals feeding on thick food has no effect on ascr#3 response.

While a great deal of work has been done to understand internal signals related to satiety and food deprivation in *C. elegans*, far less is known about how the worm’s nervous system detects and encodes the *amount* of available food. Interestingly, the TGFβ-superfamily ligand DAF-7 is important for coupling food abundance to aging and the regulation of germline progenitor proliferation [52, 53]. Our studies here indicate that *daf-7* might act upstream of *pdf-1*; understanding this will be an important area of future research. Further, PDF-2 has recently been implicated in linking nutritional status information from the intestine to locomotor state [54], raising the intriguing possibility that PDF-1 and PDF-2 communicate external and internal information about nutrition, respectively, to PDFR-1.

We find that PDFR-1 signaling does not have a single site of action with respect to the repression of ascr#3 aversion. Rather, this signal appears to act in a distributed set of interneurons to modulate motor function. Interestingly, *pdfr-1* is also known to modulate the balance between roaming and dwelling motor states, another behavior well known to be affected by food thickness [8, 40]. However, we found that restoring *pdfr-1* function to a subset of neurons that mediate these effects was unable to rescue the increased ascr#3 aversion of *pdfr-1* mutants. Future work will be required to understand how *pdfr-1* signals modulate the ascr#3 aversion circuit downstream of ASI.

How might *pdfr-1* signaling be regulated by food abundance? Because *pdf-1* appears to be more important for this function than *pdf-2*, we propose that the secretion of one or more *pdf-1*-encoded neuropeptides is regulated by a food thickness signal. Previous studies [55] as well as recent single-cell RNAseq data [47] indicate that sensory neurons are not a major site of *pdf-1* expression, indicating that sensory signals may be linked to *pdf-1* indirectly. Previous work suggests at least four distinct mechanisms that could couple the detection of food abundance to *pdf-1* activity. The physical presence of bacterial food can be detected by the deirid sensory neuron ADE [56]; dopamine released by this neuron might in turn regulate PDF-1 secretion. Bacterial food is also a rich source of chemical cues [57]; the detection of these by amphid chemosensory neurons could indirectly regulate *pdf-1*. Further, in bacteria-rich environments, the local concentrations of O_2_ and CO2 are decreased and increased, respectively, allowing worms to use the activity of gas-sensing neurons as proxy signals for food abundance [58–62]. *pdf-1* signaling could be downstream of such a signal. A final possibility has to do with the physical structure of the sparse-food environment. In our thin-food T1 conditions, bacteria are present in patches interspersed with many small food-free gaps. As animals navigate this environment, the frequency with which they cross food boundaries is markedly higher than in an abundant-food environment. Recent work has shown that worms can use this information to assess the physical structure of food patches [12]; an intriguing possibility is that this might occur at least in part via modulation of PDF-1 release and/or PDFR-1 signaling.

In males, the previously described sex-specific ability of the ADF sensory neurons to detect ascr#3 [33] overrides ASI-dependent ascr#3 aversion. Thus, in wild-type males, ablation of ASI has no effect on ascr#3 attraction. However, disabling the male-specific function of ADF by genetic feminization or ablation reveals a latent aversion to ascr#3 [33]. Here, we find that this circuit appears to be equivalent to the one that acts in hermaphrodites, in that it requires ASI and is repressed by PDFR-1 signaling. Surprisingly, however, the aversive behavior driven by this mechanism is insensitive to food thickness: even in the presence of thick food, ADF-ablated males displayed ASI-dependent avoidance of ascr#3. Thus, some aspect of the food thickness signal—either its existence or its ability to modulate the aversive signal—is hermaphrodite-specific. Interestingly, this hermaphrodite-specific feature was not eliminated when the hermaphrodite nervous system was genetically masculinized. Several possible mechanisms are consistent with this finding: the integration of food abundance and pheromone signals might require hermaphrodite-specific cues from outside the nervous system; it might require hermaphrodite-specific neurons; or it might be blocked by male-specific neurons. Further, why males lack this sensory integration is unclear. One possibility is that, rather than encoding food thickness, *pdfr-1* signaling conveys some other dimension of information in males. The important role of *pdfr-1* in promoting mate-searching behavior in males [19, 63] strongly supports this possibility.

Neuromodulation is an ancient and widespread feature of nervous systems that allows external and internal information to reconfigure the properties of neural circuits [64]. Even in simple nervous systems, myriad neuromodulatory signals exist: the *C. elegans* genome, for example, contains over 100 *flp-* and *nlp-*family neuropeptide genes, each of which can produce multiple ligands [65] (worm.peptide-gpcr.org). While significant progress has been made in determining the effects of these signals on physiology and behavior, understanding the functional significance of the modulation they bring about can be more challenging. Here, we make important progress in this area by providing evidence that PDF-1/PDFR-1 signaling serves as an internal representation of food abundance that regulates the strength of the behavioral response to stimulation by ascr#3. In *C. elegans*, *Drosophila*, and mammals, PDF-family neuropeptides (and/or the neurons that release them) have diverse functions, but in many cases, they modulate arousal and the sensitivity to external stimuli [66–69]. Our results suggest that modulation of attention to sensory stimuli could be an ancient function of this family of neuropeptides.

## METHODS

### Nematode Culture

All *C. elegans* strains were cultured using *E. coli* OP50 and NGM agar as described [70, 71]. All strains were grown at 20°C except for those containing *daf-7* mutations; these were cultured at 15°C from egg to L4 stage and then assayed at 20°C. See Table S1 for a complete list of strains used in this work. *him-5(e1490)* is considered the wild-type for these studies. All strains contained this mutation except as noted in Table S1.

### Plasmid Constructs and Transgenic Strains

To express nCre in different cells, we created a series of nCre expression plasmids. Promoters were designed from sequence information from Wormbase and were amplified from purified *C. elegans* genomic DNA using Phusion DNA polymerase (New England Biolabs). nCre expression constructs containing these promoters and the *unc-54* 3’UTR were created using Gateway cloning (Thermo Fisher Scientific). All constructs were confirmed by Sanger sequencing. See Table S2 for primer sequences.

To create conditional *pdfr-1* rescue strains, the nCre. expression plasmid was first injected into the strain UR930 with the co-injection marker Pvha-6::mCherry(mini), a gift from K. Nehrke. The resulting transgene was then crossed into the *kyEx4648* background (CX14488, generously provided by S. Flavell and C. Bargmann [40]) and maintained by picking hermaphrodites with both intestinal and pharyngeal mCherry signals.

All new extrachromosomal array transgenes were created by microinjection of DNA of interest at 20 ng/μL together with co-injection marker at 50 ng/μL. At least two lines were assayed for each new array. Genetic ablation strains were confirmed by DiI staining (for ablation of ADL, ASK, and ASI) or using a fluorescent marker (for ablation of ADF). Strains were genotyped by PCR and/or Sanger sequencing (see Table S2 for details).

### Quadrant assay with controlled bacterial thickness

Quadrant assays were performed as described previously [33] with the following modifications. To prepare lawns of controlled thicknesses, plates were seeded with bacterial cultures of defined concentration. To prepare these, 2 mL of *E. coli* OP50, freshly cultured in LB media, was collected and centrifuged (12,000 g for 30 sec). After determining the mass of the bacterial pellet, it was vigorously resuspended in sterile water (143 μL water for each 1 mg of *E. coli*) to create the T3 stock. T3 stock was diluted 0.1x to create T2 stock, which was then diluted 0.1x to create T1 stock. Each assay plate was seeded with 50 μL of stock suspension and incubated at 20°C for exactly 16 hr before t = 0 min. For three random T3 stocks, bacterial density was calculated by serial dilution to be 3.5×10^8^, 8.8×10^8^, and 2.8×10^9^ cfu/mL. After 16 hours of incubation, the lawns created with these stocks appeared identical under a stereomicroscope.

For each assay, 30 mins before t = 0 min, ten animals were picked to the center of each plate, with the experimenter blind to genotype. At t = 0, four 1 μL drops of ascr#3 or vehicle (an equivalent volume of ethanol diluted in sterile water) were dropped into each of the four quadrants. At t = 30, 60, 90, and 120 mins, the number of animals in each quadrant was scored. For each assay, the overall Quadrant Chemotaxis Index (QCI) was calculated as the mean of the four QCI values.

### Foraging assay

On the day preceding the assay, L4 hermaphrodites were picked to OP50-seeded plates (30 animals per plate) and assay plates were prepared. Guided by a custom-made transparent template, the boundaries of a center square (“source patch”) and four peripheral rectangles (“outer patches”) were drawn on the bottom of each unseeded 10-cm NGM agar plate. The shape and size of the source patch is equivalent to a single quadrant in the quadrant assay. The shape and size of each outer patch is equivalent to two adjacent quadrants. The distance from the boundary of the source patch to the nearest boundary of an outer patch is 1.8 cm. T1 or T3 bacterial stocks were made as described above. For the source patch, 12.5 μL of T1 or T3 stock suspension was dropped and carefully spread. For each outer patch, 25 μL of T3 suspension was spread. All assay plates were then incubated at 20 °C for exactly 16 hr before t = 0.

At t = 0, 10 μL ascr#3 (1 μM solution) or vehicle control (ethanol diluted in sterile water) was dropped onto the source patch under the stereomicroscope. Air bubbles were created and quickly moved around on the square food lawn to cover it completely. Once the lawn was dry (3 to 5 min), 25 WT hermaphrodites (strain DR466) and 25 ASI-ablated hermaphrodites (strain UR1110) were picked into the source patch. Each plate was incubated at 20°C for 3 hr. At t = 180 min, animals in the outer patches were counted and genotyped using GFP fluorescence.

### Statistical analysis

Unless otherwise indicated, statistical significance was assessed using a two-tailed Mann-Whitney t-test with unequal variances (to compare two genotypes) or Tukey’s multiple comparison test (to compare more than two genotypes) post one-way or two-way ANOVA, corresponding to the number of factors. Asterisks indicate *p* values associated with these tests: * *p* ≤ 0.05; ** *p* ≤ 0.005; *** *p* ≤ 0.001. For clarity, the brackets in each graph indicate all comparisons made; those with a statistically non-significant result (*p* > 0.05) are shown with dashed gray lines. For each group tested in the quadrant assay, we also carried out a one-sample t-test to ask whether there was a significant aversive or attractive response to ascr#3 (*i.e.*, to ask whether the QCI was statistically different from zero). The resulting *p* values are indicated with circles above each violin plot: ° *p* ≤ 0.05; °° *p* ≤ 0.005; °°° *p* ≤ 0.001.

## ACKNOWLEDGEMENTS

We are grateful to current and past members of the Portman lab, the University of Rochester Invertebrate Biology Group, and the Western New York Worm Group for discussion and critical feedback. We are particularly grateful to E. DiLoreto and J. Srinivasan, as well as K. Kim and P. Sengupta, for communicating unpublished results, and to F. Schroeder for providing synthetic ascr#3. We thank S. Flavell, C. Bargmann, T. Hirotsu, and D. Ferkey for generously providing transgenic strains. Some strains used in this work were provided by the *Caenorhabditis* Genetics Center, which is funded by NIH Office of Research Infrastructure Programs (P40 OD010440). These studies were funded by NIH R01 GM130136 to D.P.

## AUTHOR CONTRIBUTIONS

Conceptualization: J.L. and D.P.; Investigation: J.L.; Writing — Original Draft: J.L.; Writing — Review & Editing: D.P.; Supervision: D.P.; Funding Acquisition: D.P.

---

**Figure S1.**
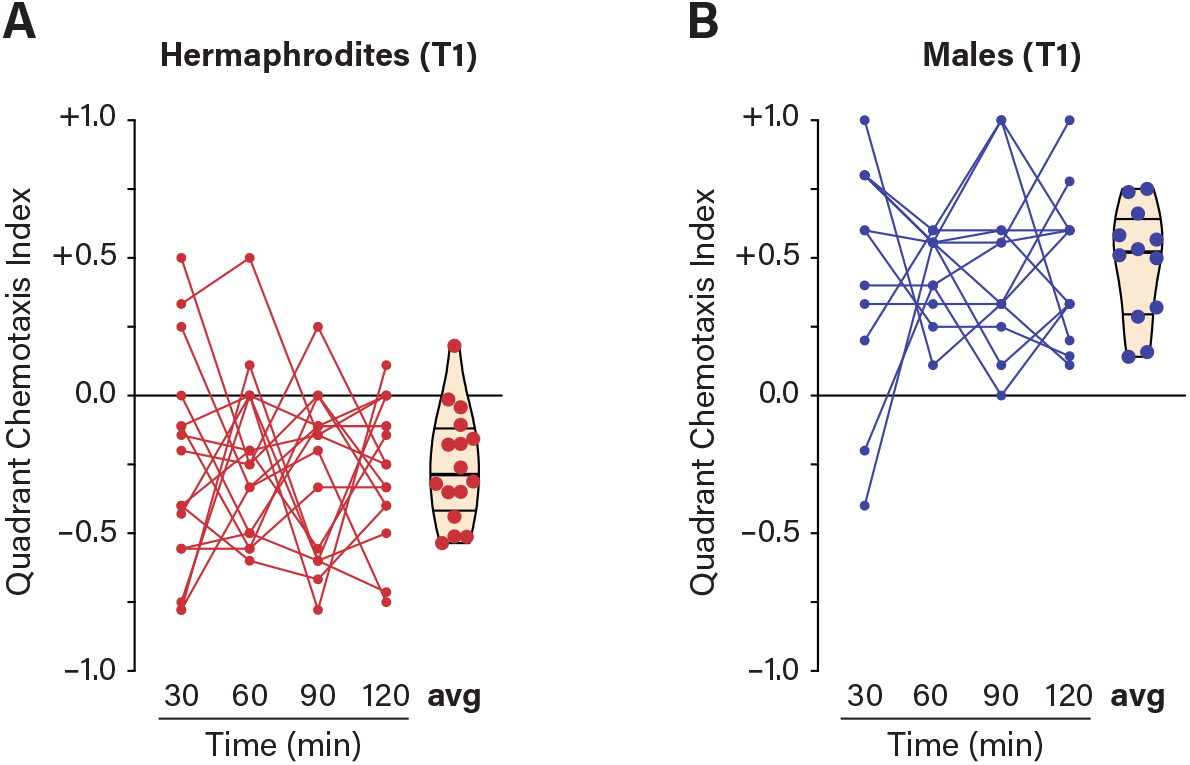
Supplementary data related to Figure 1. **(A, B)** QCIs for WT hermaphrodites (A) and males (B) in response to 1.0 μM ascr#3 with T1 food. QCI values from a given assay are connected across timepoints by a line. The violin plot at the right shows average QCI values (the mean of the four timepoints for each assay). These data are taken from Figure 1D.

**Figure S2.**
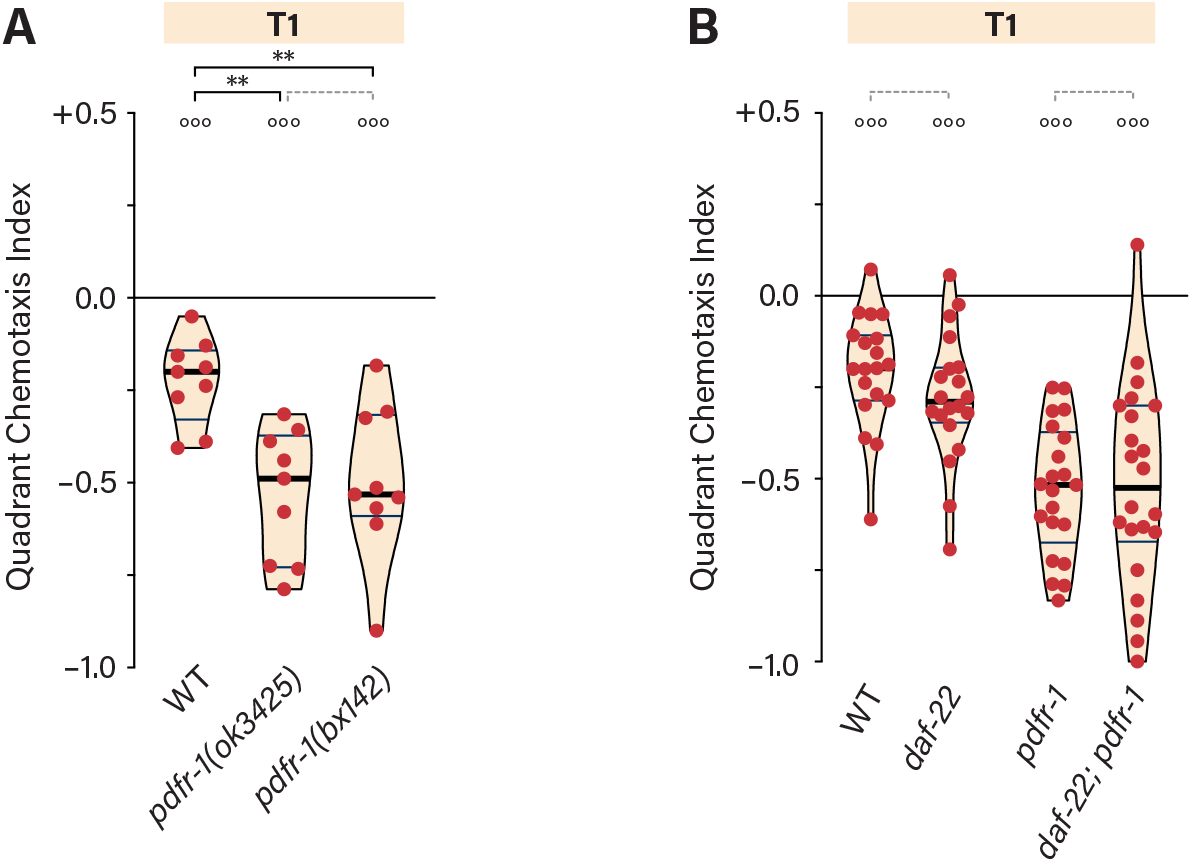
Supplementary data related to Figure 2. **(A)** QCIs for WT, *pdfr-1(ok3425)*, and *pdfr-1(bx142)* hermaphrodites with T1 food. **(B)** QCIs for WT, *daf-22*, *pdfr-1*, and *daf-22; pdfr-1* hermaphrodites with T1 food. See legend to Figure 1 for details on statistical analyses.

**Table S1.**
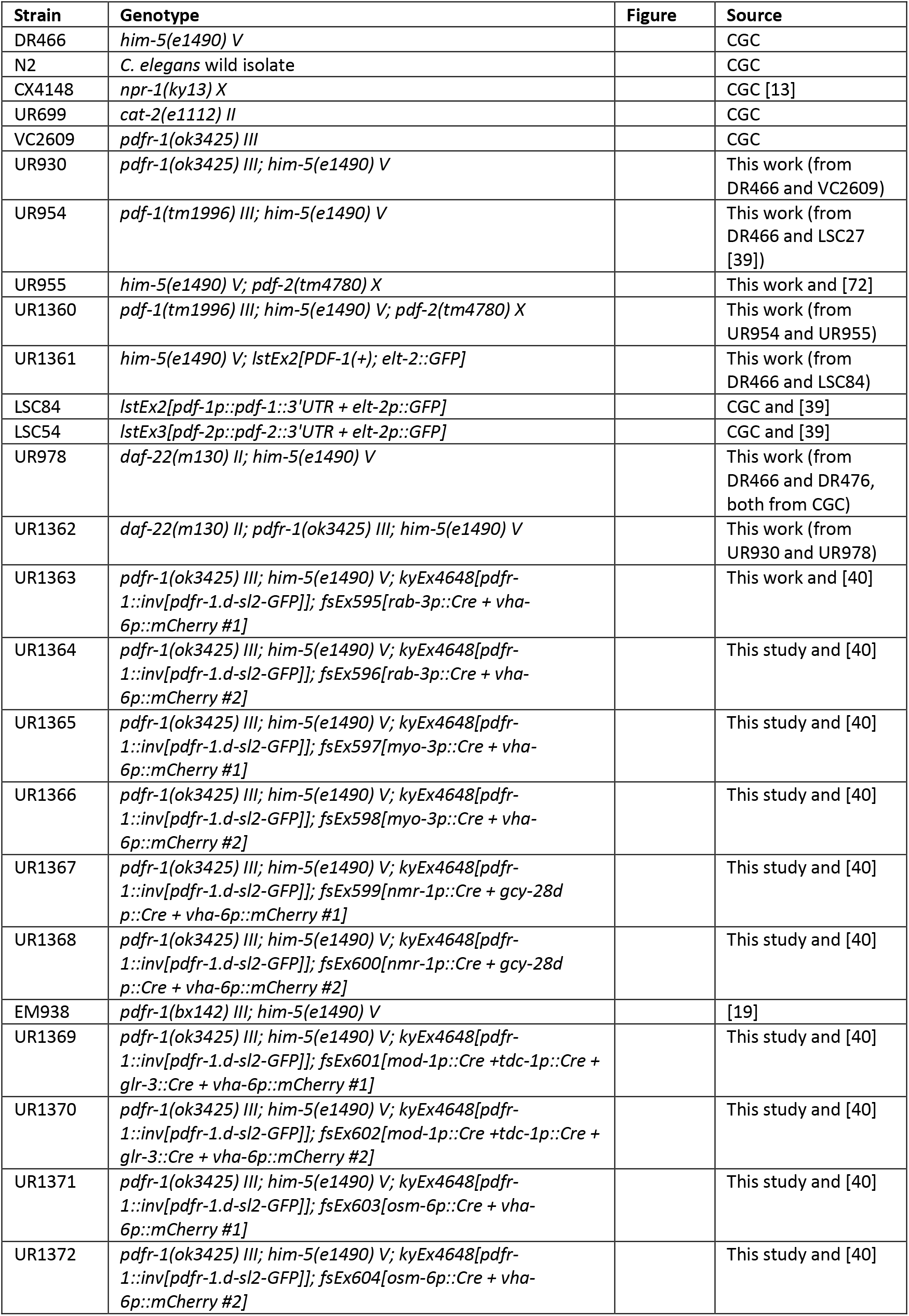
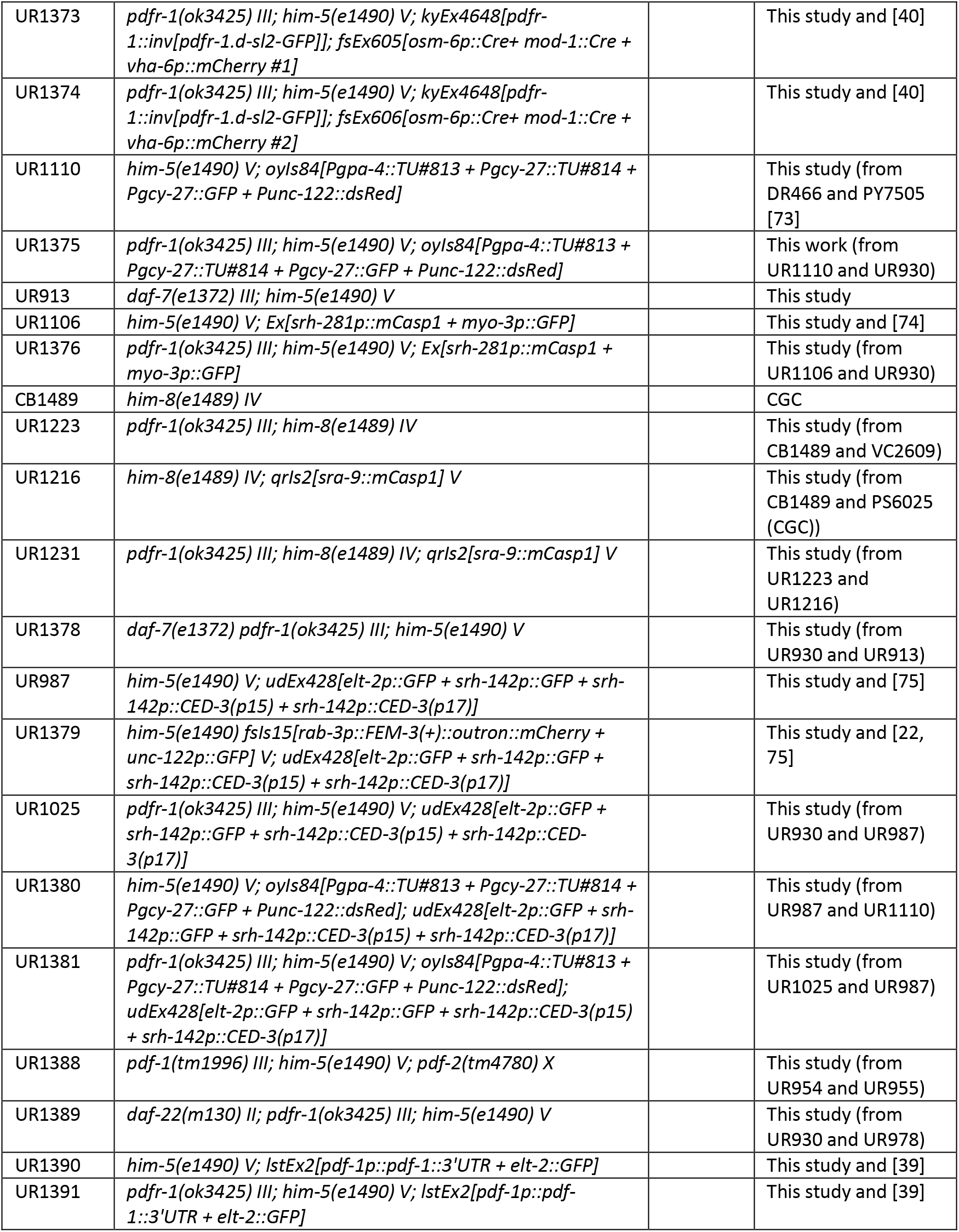
Strains used in this work.

**Table S2.**
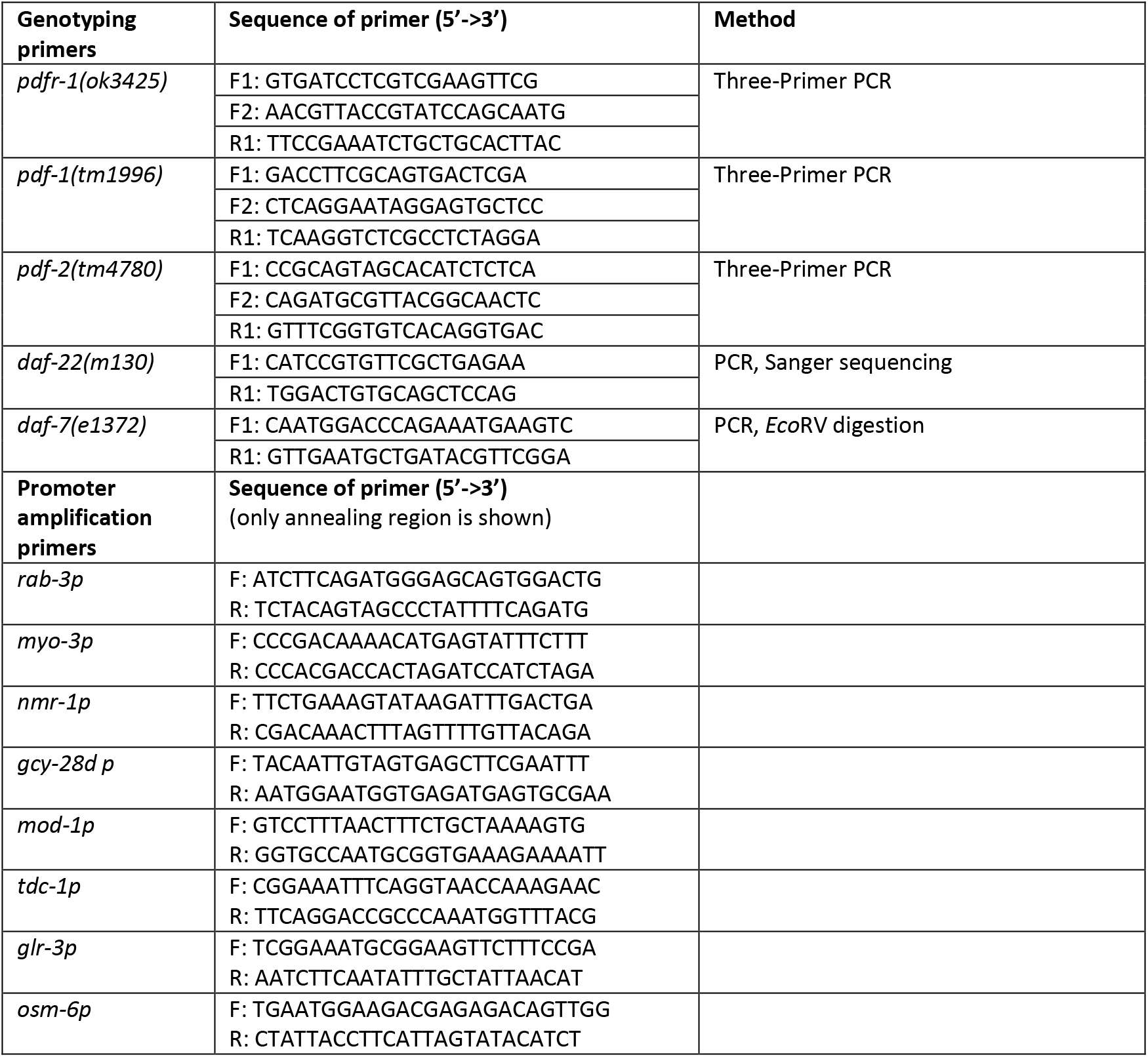
Oligonucleotide primers used in this work.

